# Structural basis for acyl group discrimination by human Gcn5L2

**DOI:** 10.1101/043364

**Authors:** Alison E. Ringel, Cynthia Wolberger

**Affiliations:** Department of Biophysics and Biophysical Chemistry, Johns Hopkins University School of Medicine, Baltimore, USA, 21205, USA; Department of Biophysics and Biophysical Chemistry, Johns Hopkins University School of Medicine, 725 N. Wolfe Street, Baltimore, USA, 21205, USA

**Keywords:** Histone acetyltransferase, acyltransferase, propionyl-CoA, butyryl-CoA

## Abstract

Gcn5 is a conserved acetyltransferase that regulates transcription by acetylating the N-terminal tails of histones. Motivated by recent studies identifying a chemically diverse array of lysine acyl modifications *in vivo*, we examined the acyl chain specificity of the acetyltransferase, human Gcn5 (Gcn5L2). Whereas Gcn5L2 robustly catalyzes lysine acetylation, the acyltransferase activity of Gcn5L2 gets progressively weaker with increasing acyl chain length. To understand how Gcn5 discriminates between different acyl-CoA molecules, we determined structures of the catalytic domain of human Gcn5L2 bound to propionyl-CoA and butyryl-CoA. Although the active site of Gcn5L2 can accommodate propionyl-CoA and butyryl-CoA without major structural rearrangements, butyryl-CoA adopts a conformation incompatible with catalysis that obstructs the path of the incoming lysine residue and acts as a competitive inhibitor for Gcn5L2 versus acetyl-CoA. These structures demonstrate how Gcn5L2 discriminates between acyl chain donors and explain why Gcn5L2 has weak activity for acyl moieties that are larger than an acetyl group.

## 1. Introduction

Lysine acetylation is an abundant post-translational modification (Weinert *et al*., 2011; Choudhary *et al*., 2009) that changes the overall size and charge of the modified residue. Several classes of enzymes are known to catalyze site-specific lysine acetylation (Roth *et al*., 2001; Yang, 2004; Marmorstein & Trievel, 2009), many of which localize to the nucleus and modify lysine residues on histones (Lee & Workman, 2007). These enzymes are collectively referred to as either <u>***H***</u>istone <u>***A***</u>cetyl<u>***T***</u>ransferases (HATs) or Lysine (<u>***K***</u>) <u>***A***</u>cetyl<u>***T***</u>ransferases (KATs), the latter to reflect their ability to acetylate non-histone substrates (Glozak *et al*., 2005). KATs are divided into several main families based on both structural homology and the presence of sequence conservation within their catalytic domains (Marmorstein & Trievel, 2009). Although different KAT families employ distinct kinetic mechanisms to catalyze acetyl transfer, they all share a common dependence on the nucleotide cofactor, acetyl-CoA, as an acetyl donor (Berndsen & Denu, 2008).

Gcn5 is a member of the GNAT (<u>***G***</u>cn5-related-<u>***N***</u>-<u>***A***</u>cetyl<u>***T***</u>ransferase) family of histone acetyltransferases that acetylates the N-terminal tails of histones H3 and H2B at the promoters of inducible genes (Grant *et al*., 1997) and is broadly implicated in transcriptional regulation (Huisinga & Pugh, 2004). The kinetic mechanism of Gcn5 has been studied extensively (Tanner *et al*., 1999; Tanner, Langer, Kim, *et al*., 2000) and the Gcn5 catalytic domain from several organisms has been crystallized in the presence of various combinations of substrates (Roth *et al*., 2001; Poux *et al*., 2002). The active site of Gcn5 contains two grooves where acetyl-CoA and peptide bind, which intersect near the β-mercaptoethylamine moiety of coenzyme A and the target lysine (Rojas *et al*., 1999; Poux *et al*., 2002). The ternary complex between Gcn5, acetyl-CoA, and peptide forms through a fully ordered mechanism (Tanner, Langer, Kim, *et al*., 2000), as binding to acetyl-CoA brings about a structural rearrangement that widens the peptide-binding groove within the Gcn5 active site (Clements *et al*., 1999; Trievel *et al*., 1999; Rojas *et al*., 1999; Lin *et al*., 1999). Because Gcn5 primarily recognizes features of the CoA pantetheine arm and not the acetyl group (Poux *et al*., 2002; Clements *et al*., 1999; Rojas *et al*., 1999; Lin *et al*., 1999), Gcn5 binds with similar affinity to acetyl-CoA and free CoA (Tanner, Langer, Kim, *et al*., 2000). Whether the active site of Gcn5 can accommodate other kinds of CoA molecules with bulkier acyl chains, however, is not known.

Recent studies have found that lysine residues are modified by a chemically diverse array of acyl chains *in vivo* (Lee, 2013), raising the possibility that some KATs might be able to utilize acyl-CoA cofactors other than acetyl-CoA to catalyze lysine acylation. Members of three different KAT families catalyze lysine acylation *in vitro*: p300/CBP catalyzes propionylation, butyrylation, and crotonylation of histones and p53 (Chen *et al*., 2007; Sabari *et al*., 2015), yeast Esa1, a member of the MYST family of acetyltransferases, catalyzes propionylation of histone H4 peptides (Berndsen *et al*., 2007), and human P/CAF, which is closely related by sequence homology to Gcn5, catalyzes propionylation of histone H3 peptides (Leemhuis *et al*., 2008). Some KATs are clearly promiscuous with regards to acyl chain identity, but the mechanisms employed by acyltransferases to discriminate between different acyl-CoA molecules are still largely unknown.

In this study, we characterize the acyl chain specificity of human Gcn5, which catalyzes acetylation of histone peptides much more quickly than either propionylation or butyrylation. To understand how Gcn5 discriminates between different acyl chain donors, we determined structures of the catalytic domain of human Gcn5L2 in complex with propionyl-CoA and butyryl-CoA to 2.0 and 2.1 Å resolution, respectively. These structures reveal that the active site of Gcn5 can accommodate longer acyl chains without major structural rearrangements, but that the butyryl chain would sterically clash with an incoming lysine residue. Consistent with this active site architecture, we show that butyryl-CoA acts as a competitive inhibitor versus acetyl-CoA for human Gcn5L2. These findings raise the possibility that some acyl-CoA molecules might function as natural inhibitors of Gcn5 *in vivo*, and have important implications for the regulation of KATs in response to metabolic changes.

## 2. Methods

### 2.1 Protein expression and purification

A plasmid encoding the his-tagged catalytic domain of human Gcn5L2 (hsGcn5L2) under T7-induction was obtained from Addgene (Plasmid No.: 25482). The protein was expressed and purified as previously described (Schuetz *et al*., 2007). Purified protein was dialyzed into 20 mM HEPES, pH 7.5, 150 mM NaCl, and 1 mM DTT, concentrated to 9 mg/mL, flash-frozen in liquid nitrogen, and stored at ‐80°C.

### 2.2 Enzymatic Assays

Kinetic measurements comparing rates of acetylation, propionylation and butyrylation were performed using the 5,5’-dithiobis-(2-nitrobenzoic acid) (DTNB) assay (Berndsen *et al*., 2007) with the following modifications. Reactions contained 10 μM hsGcn5L2 catalytic domain, 250 µM histone H3 peptide aa1-21 (purchased from United Peptide at >90% purity), 100 mM HEPES, pH 7.6, 50 mM NaCl, and 500 µM acetyl-CoA (Sigma No.: A2181), propionyl-CoA (Sigma No.: P5397) or butyryl-CoA (Sigma No.: B1508). Each reaction was incubated for five minutes at 37°C before adding acyl-CoA, and then maintained at 37°C for the remainder of the experiment. Initially, six data points were collected to find a time frame where acyl-CoA consumption was linear with time. The reaction was quenched at the indicated time points by the addition of two volumes of quenching buffer (3.2 M guanidine-HCl and 100 mM sodium phosphate pH 6.8). After all the samples had been collected, one volume of 4 mM DTNB (Sigma No.: D218200) dissolved in 100 mM sodium phosphate, pH 6.8 was added. Samples were moved to a 384-well polystyrene clear-bottom plate (Grenier Bio-One) and absorbance at 412 nM was measured in a POLARstar Omega plate reader (BMG Labtech). Absorbances were converted to concentrations using a standard curve generated by reacting increasing concentrations of CoA (Sigma No.: C3019) with DTNB using an extinction coefficient for 3-thio-6-nitrobenzoate (TNB) of ε_412nm_ = 13,700 M^-1^ cm^-1^. Subsequent reactions were performed in triplicate and quenched after 0.5 minutes (acetyl-CoA), 5 minutes (propionyl-CoA), or 20 minutes (butyryl-CoA). All acylation rates were corrected by subtracting the rate of acyl-CoA consumption by Gcn5L2 in the absence of peptide.

Steady-state kinetic titrations varying the acetyl-CoA or butyryl-CoA concentration were performed with a continuous spectrophotometric assay as previously described (Berndsen & Denu, 2005). Briefly, the acetyl-CoA or butyryl-CoA concentration was varied between 0.25 µM and 100 µM in the presence of 50 nM hsGcn5L2 and 300 µM histone H3 peptide. Reactions were performed in a total volume of 50 µL at 37°C in 384-well plates (Greiner Bio-One) and initiated with acyl-CoA. The absorbance at 340 nm was monitored continuously using a POLARstar Omega plate reader (BMG Labtech) for 5-20 minutes, and converted into the molar concentration of NADH using Beer’s Law, assuming ε_340nm_ = 6220 M^-1^ cm^-1^. As controls, rate measurements were performed at each concentration of acyl-CoA in the absence of peptide. Each measurement was performed in triplicate, and reaction velocities in the presence of peptide were blanked by the rate of reaction in the absence of peptide. Blanked rates were normalized to enzyme concentration, plotted as a function of substrate concentration, and fit to the Michaelis-Menten equation using non-linear least squares regression in GraphPad Prism 5. Butyryl-CoA inhibition measurements were also performed with the enzyme-coupled assay. Reaction velocities were measured in the presence of 0.5 µM to 10 µM acetyl-CoA and 50 nM hsGcn5L2 with increasing concentrations of butyryl-CoA (0, 50, 100, or 300 µM). Under these conditions, consumption of butyryl-CoA by hsGcn5L2 is undetectable by the same assay. Blanked rates were normalized to enzyme concentration and the resulting curves were globally fit to a competitive inhibition model in Graphpad Prism 5.

### 2.3 HAT Domain Crystallization

Propionyl-CoA and butyryl-CoA were diluted in 20 mM HEPES, pH 7.5 and stored at ‐20°C at a concentration of 20 mM calculated using ε_260nm_ = 16,400 M^-1^ cm^-1^. Purified human Gcn5L2 aa497-662 was mixed with each acyl-CoA to a final concentration of 1.6 mM acyl-CoA and 7.9 mg/mL protein. NaCl was added to a final concentration of 125 mM from a 5 M stock, and the resulting mixture was incubated on ice for 30 minutes. Both complexes were crystallized using hanging drop vapor diffusion by mixing 1 µL of protein: acyl-CoA complex with 1 µL of well solution. Human Gcn5L2 bound to propionyl-CoA was crystallized in 20% (v/v) ethanol and 100 mM Tris, pH 9.0. Human Gcn5L2 bound to butyryl-CoA was crystallized in 10% (v/v) 2-propanol, 3% glycerol, 100 mM HEPES, pH 7.8, and 11% (w/v) PEG 4,000. Crystals were cryoprotected by soaking in well solution supplemented with 9% sucrose, 4% glucose, 8% ethylene glycol, and 8% glycerol. Prior to data collection, crystals were flash frozen in a liquid nitrogen stream.

### 2.4 Data Collection and Processing

Diffraction data were collected using a Rigaku FR-E SuperBright x-ray generator at a wavelength of 1.54 Å and recorded with a Saturn 944+ CCD detector. Data were processed with HKL2000 (Otwinowski & Minor, 1997). The structures were solved using molecular replacement with MOLREP from the CCP4 suite using the coordinates of human Gcn5L2 (PDB ID 1Z4R) as a search model (Vagin & Teplyakov, 2010; Vagin & Teplyakov, 1997). Refinement was carried out using REFMAC5 from the CCP4 suite (Winn *et al*., 2011; Murshudov *et al*., 1997) and the graphics program COOT for model-building (Emsley & Cowtan, 2004). Simulated annealing omit maps were generated by removing either propionyl‐ or butyryl-CoA from the refined structures, fitting acetyl-CoA into the ligand density, and performing three rounds of refinement with Phenix including two cycles of simulated annealing (Adams *et al*., 2010). Data collection and refinement statistics are shown in Table 1. RMSD calculations were performed using PDBefold from the EMBL-EBI website. Structure figures were generated with Pymol Molecular Graphics System, Version 1.7.4 Schrödinger, LLC.

**Table 1.**
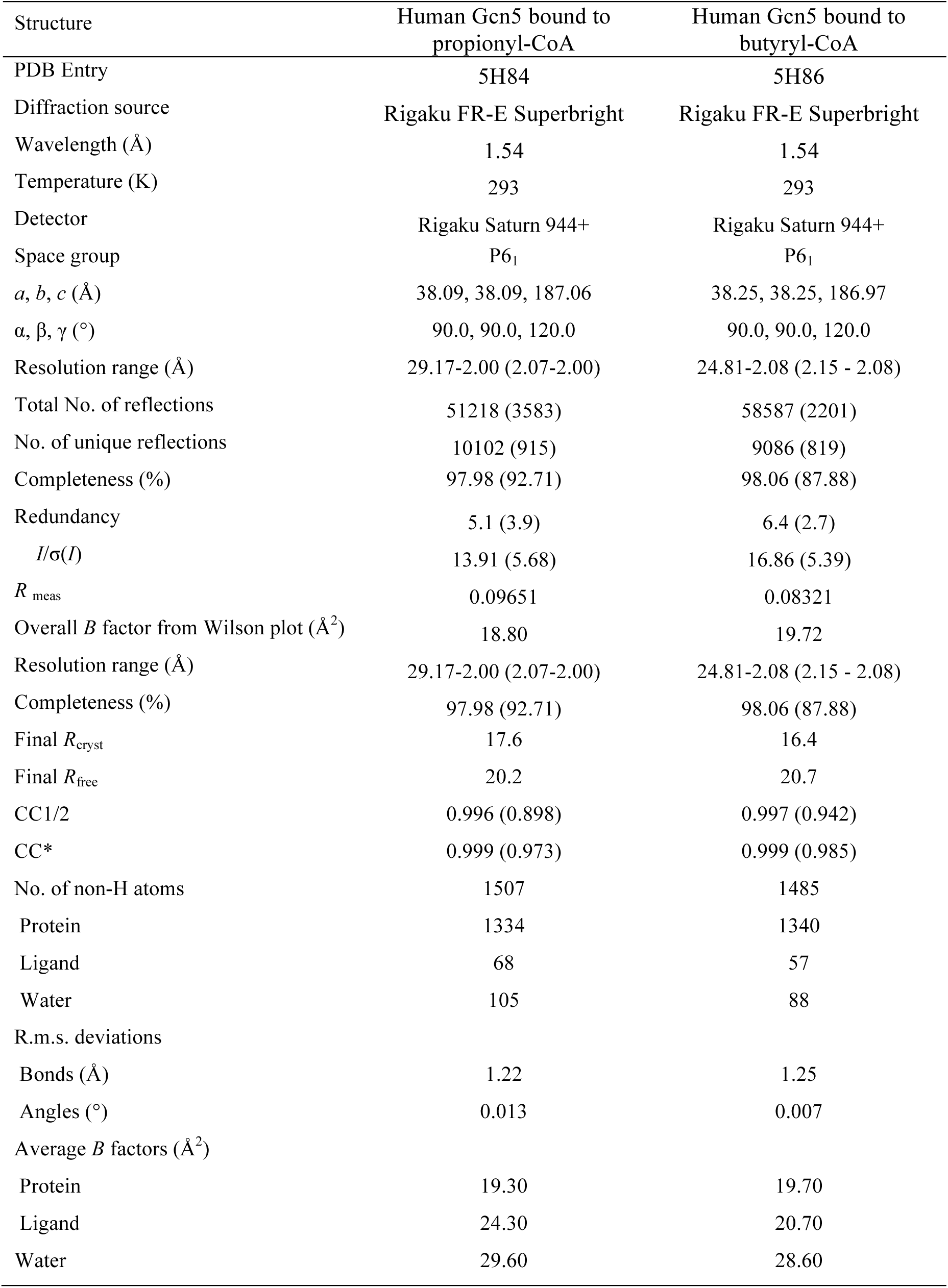
Data collection, refinement, and model statistics Values for the outer shell are given in parentheses.

### 2.5 PDB accession codes

Structures and amplitudes have been deposited in the Protein Data Bank with accession codes 5H84 (propionyl-CoA) and 5H86 (butyryl-CoA).

## 3. Results

### 3.1 Gcn5 is a weak acyltransferase

Previous studies of the P/CAF acetyltransferase, whose catalytic domain shares 95% sequence identity with Gcn5, showed that P/CAF catalyzes histone propionylation with similar kinetics to acetylation (Leemhuis *et al*., 2008). Compared to acetyl-CoA, the P/CAF *K*_m_ for propionyl-CoA is only fourfold weaker, corresponding to a six-fold decrease in catalytic efficiency (Leemhuis *et al*., 2008). To determine whether human Gcn5 can similarly use other acyl-CoAs as a cofactor, we measured Gcn5 activity in the presence of either propionyl-CoA or butyryl-CoA and histone H3 peptide. We found that human Gcn5L2 efficiently acetylates and propionylates peptides, while its butyrylating activity is nearly undetectable (Figure 1). Under these experimental conditions, Gcn5L2 propionylates histone peptides approximately nine-fold more slowly and butyrylates peptides nearly 400-fold more slowly compared to its acetyltransferase activity (Figure 1). Based on these relative rate measurements, Gcn5L2 is unlikely to contribute significantly to lysine butyrylation *in vivo* but may be capable of catalyzing lysine propionylation under physiological conditions.

**Figure 1.**
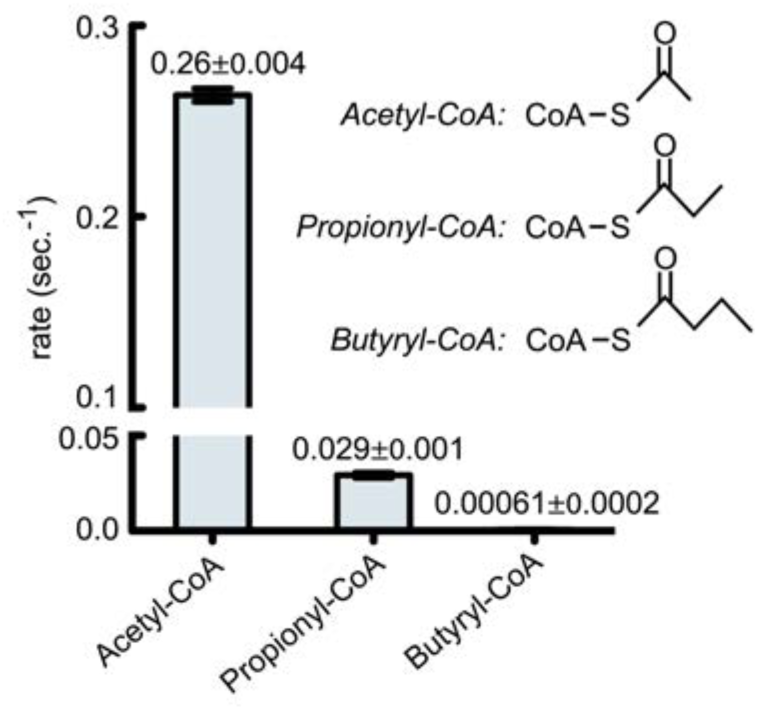
Acyltransferase activity of hsGcn5L2. Rate of catalysis by hsGcn5L2 (10 μM) were measured using different acyl-CoA cofactors (500 μM) and N-terminal histone H3 peptide (250 μM) containing the sequence: ARTKQTARKSTGGKAPRKQLA.

### 3.2 Structures of hsGcn5L2 bound to propionyl-CoA and butyryl-CoA

To elucidate the structural basis for the ability of Gcn5L2 to discriminate among different acyl-CoA cofactors, we determined the structure of human Gcn5L2 bound to propionyl-CoA and butyryl-CoA to 2.0 Å and 2.1 Å resolution, respectively. Refinement statistics for each structure are summarized in Table 1. Simulated annealing omit maps show clear density corresponding to the extra methyl group for propionyl-CoA (Figure 2A) or extra ethyl chain for butyryl-CoA (Figure 2B). Compared to the structure of human Gcn5L2 bound to acetyl-CoA (Schuetz *et al*., 2007), the structures reported here are very similar; the root-mean-square difference (RMSD) in Cα positions is 0.13 Å for the structure of human Gcn5L2 in complex with propionyl-CoA and 0.24 Å for its structure in complex with butyryl-CoA.

**Figure 2.**
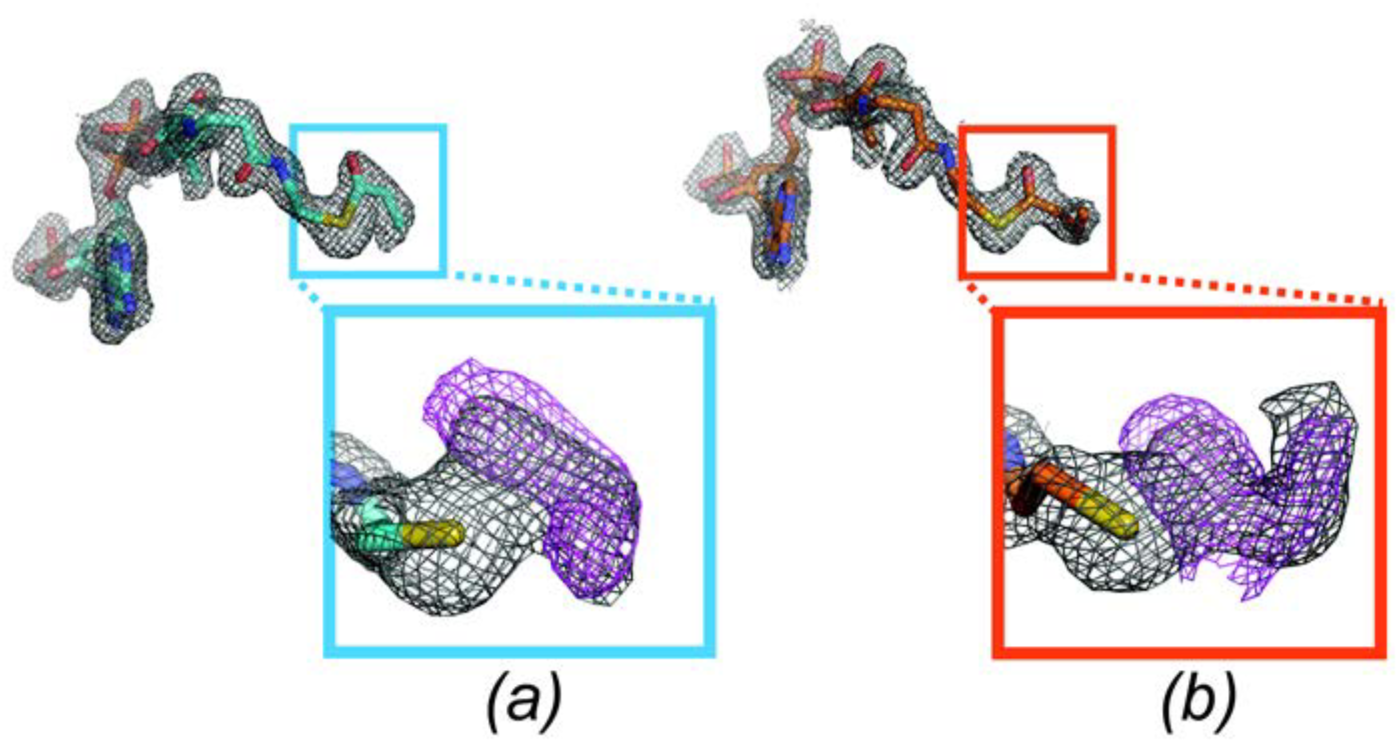
Simulated annealing omit maps demonstrate clear density for the acyl cofactors. Omit maps were calculated with either the (A) propionyl or (B) butyryl moieties removed. 2Fo-Fc maps are contoured at 1σ (gray) and simulated annealing omit maps are contoured at 2.5-3σ (magenta).

The active site of Gcn5 contains three features that facilitate transfer of the acyl chain to lysine: an active site glutamate that functions as a general base (Tanner *et al*., 1999), a structurally conserved water molecule that forms a proton wire between the general base and incoming lysine (Rojas *et al*., 1999), and residues that stabilize the position of the acyl-CoA (Figure 3A & 3B) (Rojas *et al*., 1999; Schuetz *et al*., 2007). This active site geometry is preserved in the two acyl-CoA bound structures reported here, including the orientation of the acyl-CoA thioester, which is coordinated by the backbone amide of cysteine 579, and the position of the water molecule, which is hydrogen bonded to glutamate 575 (Figure 3C & 3D). The conservation of active site geometry rules out the possibility that butyryl-CoA binding slows down Gcn5 catalytic activity by misaligning active site residues. The pantethine arm and adenine moieties of coenzyme A superimpose well between all three acyl-CoA molecules; what differs is the respective position of the acetyl, propionyl, and butyryl chains (Figure 3E). Although the positions of the C2 carbons in all three acyl-CoA molecules are the same (Figure 3F, 3G, & 3H), the torsion angle formed between the sulfur-C1 and C2-C3 bonds in propionyl-CoA is 24.5° (Figure 3I), compared to ‐61° for butyryl-CoA (Figure 3J). Whereas the C3 carbon in propionyl-CoA fits within the active site cleft of human Gcn5 (Figure 3C), butyryl-CoA binds in an orientation that places the terminal methyl group (C4) facing the solvent, since the catalytic water molecule blocks it from occupying the Gcn5 active site cleft (Figure 3D). Although longer acyl chains could also bind in this orientation, we note that molecules like crotonyl-CoA, which contain unsaturated carbon-carbon bonds that do not freely rotate, cannot adopt conformations compatible with this geometry.

**Figure 3.**
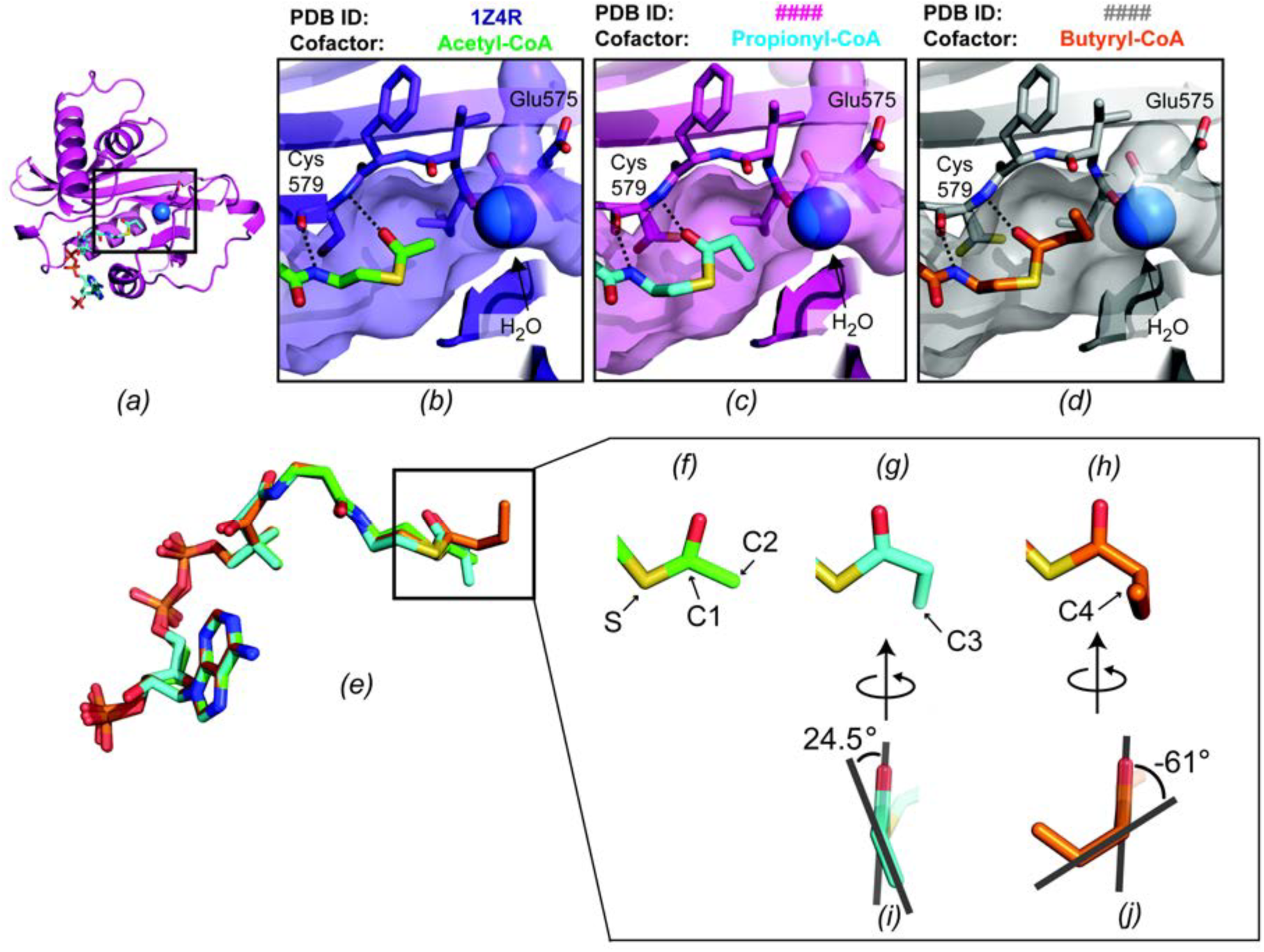
The position of the acyl chain varies in the Gcn5 active site. (A) Overall structure of the catalytic domain of hsGcn5L2, shown in cartoon representation, bound to propionyl-CoA (cyan). The catalytic site is outlined with a black rectangle, where the catalytic water molecule is shown as a blue sphere and the acyl-CoA is depicted in stick representation. Close-up views of the active site of hsGcn5L2 bound to: (B) acetyl-CoA (green), (C) propionyl-CoA (cyan), or (D) butyryl-CoA (orange). (E) Structural alignment of the three acyl-CoA molecules in the Gcn5 active site. Close-up views of the different acyl groups: (F) acetyl-CoA, (G) propionyl-CoA, and (H) butyryl-CoA. Torsion angle adopted by the (I) propionyl and (J) butyryl moieties.

### 3.3 Model of the ternary complex with peptide and CoA

Since Gcn5 uses a direct-transfer mechanism to catalyze lysine acetylation (Tanner, Langer, Kim, *et al*., 2000), Gcn5 must bind to the acyl-CoA cofactor and the peptide substrate at the same time. To determine whether the conformations adopted by propionyl-CoA and butyryl-CoA in complex with Gcn5L2 are compatible with peptide binding, models of human Gcn5L2 bound to each acyl-CoA molecule and a histone peptide (Figure 4A) were generated based on the structure of Tetrahymena Gcn5 bound to a bisubstrate analog consisting of CoA covalently linked to a histone peptide (PDB ID 1M1D) (Poux *et al*., 2002). In our model of human Gcn5L2 bound to acetyl-CoA and peptide, the incoming lysine residue makes an angle of 105° with the acetyl thioester (Figure 4B), which is a reasonable angle of attack for a carbonyl group by a nucleophile (Burgi *et al*., 1974). Propionyl-CoA also adopts a conformation compatible with this angle of attack, as the position of the terminal methyl is in the same plane as the thioester, which leaves the lysine attack trajectory open (Figure 4C). By contrast, the terminal methyl of butyryl-CoA projects into the channel occupied by the lysine (Figure 4D). The Gcn5L2 active site cannot accommodate butyryl-CoA without ejecting the catalytic water molecule, so the butyryl chain sterically clashes with the incoming lysine. This explains why Gcn5L2 is a poor butyryltransferase, as the acyl chain, catalytic water molecule (Figure 3C), and incoming lysine (Figure 4D) cannot all fit into its active site.

**Figure 4.**
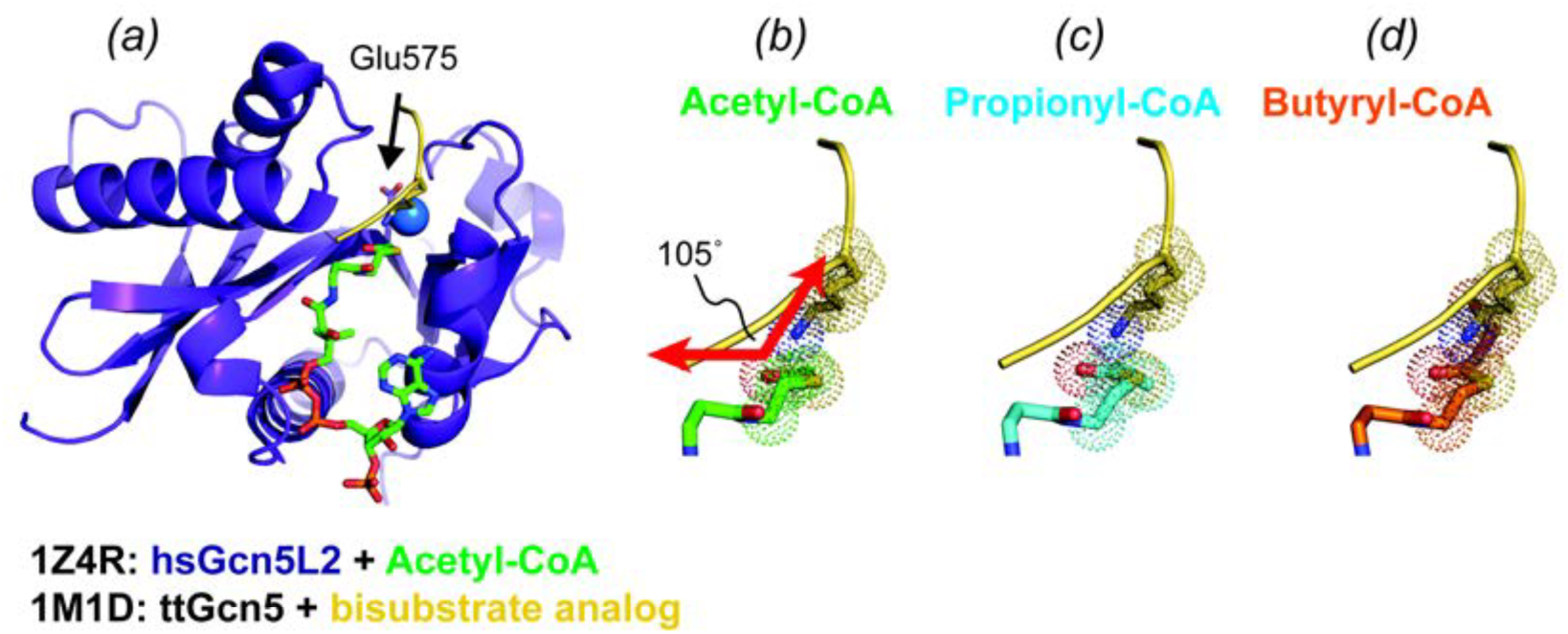
Model of the ternary complex between hsGcn5L2, different acyl-CoA molecules, and peptide. (A) Overall model with hsGn5L2 colored purple, the peptide colored yellow, and acetyl-CoA colored green. The catalytic water molecule is depicted as a blue sphere. Close-up views of the arrangement between the incoming lysine (yellow) and (B) acetyl-CoA shown in green, (C) propionyl-CoA shown in cyan, or (D) butyryl-CoA shown in orange.

### 3.4 Butyryl-CoA is a competitive inhibitor of acetylation by human Gcn5

Our results suggest that a naturally occurring acyl-CoA molecule, such as butyryl-CoA, could inhibit Gcn5 activity by binding to the enzyme in a way that prevents lysine from entering its active site. Since butyryl-CoA is a poor substrate for Gcn5 (Figure 1) but is still able to bind in the active site (Figure 3D), we wondered whether it might act as a competitive inhibitor versus acetyl-CoA. To test this idea, we measured acetylation rates as a function of acetyl-CoA concentration in the presence of increasing concentrations of butyryl-CoA. As shown in Figure 5A, Gcn5 is robustly acetylates histone peptides under the same conditions where butyrylation is nearly undetectable. Fitting our initial velocity measurements to the Michaelis-Menten equation, we determine a K_*m*_ for acetyl-CoA of 0.91 ± 0.09 µM (Figure 5A) which is comparable to previously reported K_*m*_ values for yeast Gcn5, human Gcn5, and human P/CAF (Poux *et al*., 2002; Tanner, Langer, Kim, *et al*., 2000; Tanner, Langer & Denu, 2000; Langer *et al*., 2002). We next measured acetylation kinetics in the presence of increasing concentrations of butyryl-CoA and fit the resulting curves to either competitive, noncompetitive, or uncompetitive inhibition models. Competitive inhibition clearly fits the data the best, as the sum of the squares of the residuals normalized to the degrees of freedom is 0.053 for the competitive model, compared to 0.24 and 0.26 for noncompetitive and uncompetitive inhibition, respectively. We measure an inhibition constant (*K*_*i*_) of 5.6 ± 0.7 µM from our global fit to a competitive inhibition model (Figure 5B). Taken together with our structural findings (Figure 4A & 4D), these data indicate that butyryl-CoA competitively inhibits acetylation by Gcn5 by binding to the free form of the enzyme and preventing acyl chain transfer.

**Figure 5.**
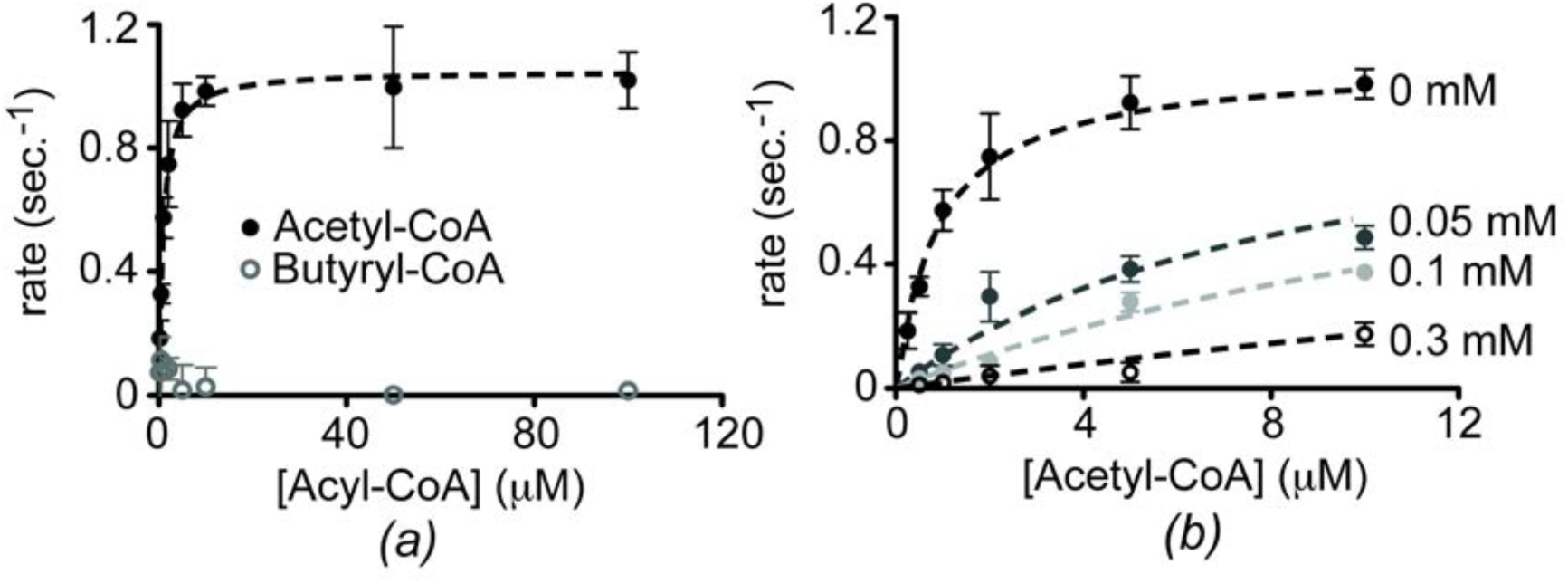
Butyryl-CoA competitively inhibits acetylation by human Gcn5. (A) Steady-state kinetic analysis comparing acetylation and butyrylation by human Gcn5 using a fixed concentration of 300 μM H3 peptide. By fitting this curve to the Michaelis-Menten equation, we find *K*_m_ = 1.05 ± 0.02 μM and *k*_*cat*_ = 0.91 ± 0.09 sec^-1^. (B) Acetylation kinetics monitored in the presence of increasing concentrations of butyryl-CoA (0-0.3 mM). Data were globally fit to a competitive inhibition model.

## 4. Discussion

We have determined crystal structures that describe how Gcn5 accommodates propionyl-CoA in its active site and provide a structural mechanism that explains our biochemical data showing that human Gcn5 discriminates between different acyl-CoA molecules. Since unsaturated acyl chains greater than three carbons in length (propyl groups) cannot fit into the Gcn5 active site, butyryl-CoA binds in a conformation that is incompatible with catalysis. The butyryl-CoA C3 and C4 carbons occupy the channel for the incoming lysine (Figure D), which prevents the peptide substrate from accessing the active site cleft of Gcn5. We further show that butyryl-CoA is a competitive inhibitor versus acetyl-CoA for human Gcn5 (Figure 5C), raising the question whether fluctuating levels of acyl-CoA molecules in cells may regulate the activity of Gcn5.

Coenzyme A is a common nucleotide cofactor that carries many different kinds of acyl groups *in vivo* (King & Reiss, 1985) and many metabolic processes produce or consume acyl-CoAs (Albaugh *et al*., 2011). As a result, intracellular concentrations of different acyl-CoA species change in response to metabolic fluctuations (Hosokawa *et al*., 1986; Palladino *et al*., 2012; King & Reiss, 1985). For example, measurements of the intracellular concentration of acetyl-CoA vary based on nutrient availability, and range from 3 to 30 µM in yeast (Cai *et al*., 2011; Weinert, 2014) and 2 to 13 μM in human cells (Lee *et al*., 2014). With a *K*_m_ for acetyl-CoA of 0.91 ± 0.09 μM (Figure 5A), acetyl-CoA availability may regulate the activity of human Gcn5 (Albaugh *et al*., 2011). Consistent with this, Gcn5-catalyzed histone acetylation is induced under growth conditions with high intracellular levels of acetyl-CoA (Cai *et al*., 2011). It is not yet known whether the intracellular concentrations of other acyl-CoA species are high enough to impact acetylation by KATs like Gcn5. Although studies quantifying absolute concentrations of propionyl-CoA and butyryl-CoA in cells have not been done, measurements of the relative abundance of different acyl-CoAs in fasting rat (King & Reiss, 1985) and mouse (Palladino *et al*., 2012) livers found roughly 4:2:1 molar ratios of acetyl, propionyl, and butyryl-CoA. With an inhibition constant of 5.6 ± 0.7 μM for butyryl-CoA, it is possible that intracellular acyl-CoA ratios could regulate the activity of Gcn5. As a result, the activity of Gcn5 would be sensitive to metabolic flux, as the relative amounts of different acyl-CoA species change in response to metabolic activity (Hosokawa *et al*., 1986; Palladino *et al*., 2012; King & Reiss, 1985).

Other CoA-based molecules have been implicated as acetyltransferase inhibitors *in vivo* and *in vitro*. Free CoA is a potent competitive inhibitor of yeast Gcn5 (Tanner, Langer, Kim, *et al*., 2000) and human P/CAF (Tanner, Langer & Denu, 2000) *in vitro*, with inhibition constants (*K*_*i*_) of 6.7 μM and 0.44 μM, respectively. Combined with the observation that free CoA is present at roughly equimolar concentrations to acetyl-CoA in cells (Gao *et al*., 2007; Lee *et al*., 2014), it is plausible that the ratio of acetyl-CoA to CoA may modulate KAT activity (Albaugh *et al*., 2011). Interestingly, a recent study profiling the acyl chain specificity of Gcn5 observed potent inhibition by long fatty acyl-CoA molecules like palmitoyl-CoA (Montgomery *et al*.), further supporting the idea that acyl-CoA molecules may function as natural KAT inhibitors. Although relatively few synthetic inhibitors for KATs have been developed, some of the most potent compounds exploit CoA-based scaffolds with structural complementarity to the active site (Furdas *et al*., 2012). Bisubstrate analogues comprised of CoA-peptide conjugates mimic the ternary complex and inhibit Gcn5 at micromolar concentrations (Poux *et al*., 2002). In light of these observations, the structures presented here suggest that unsaturated acyl-CoAs may well act as natural acetyltransferase inhibitors *in vivo* and provide a clue as to how other CoA-based scaffolds may be exploited to design future generations of acetyltransferase inhibitors.

## Acknowledgements

This work was supported by a grant from the National Institute of General Medical Sciences, GM098522.

